# Predicting the distribution of common wild mammal species across Europe – are there sufficient occurrence data?

**DOI:** 10.1101/2025.02.27.640497

**Authors:** S Croft, D Warren, JA Blanco-Aguiar, GC Smith

## Abstract

**Context:** Knowledge of where wildlife species are and in what number is critical to support robust contingency planning for diseases affecting animal and human health. For common widespread species in particular, comprehensive surveillance is impractical and challenging to coordinate. This is where models can be useful, to fill in the gaps and direct survey efforts to maximise understanding. The ENETWILD project was commissioned by EFSA in 2017 to review available data and modelling methods to predict distributions for key species involved in diseases of concern (e.g., wild boar and African swine fever). Here, we outline the latest methodology to predict species distributions based on occurrence data and outline the areas where further data would be most helpful.

**Method:** We present a generic framework to model the distribution of common mammal species in Europe, using global data. Based on occurrence records from large scale repositories, our method first attempts to address known biases to produce a presence-absence dataset and then applies random forest modelling to predict habitat suitability. We apply this approach to twelve species spanning distinct survey methodologies within mammal recording (visual observation, trapping, and bats/audio monitoring).

**Results:** Model performance was acceptable across all species within the limits of the available training data (AUC close to or above 0.7). However, beyond these limits the reliability of prediction was substantially reduced. Assessment of the available data suggested large areas of Eastern and Southern (particularly mountainous) parts of Europe have a distinct environmental signature not sufficiently captured by the existing sampling.

**Conclusions:** Within Europe there remain environmental conditions which are not well represented by existing surveillance, predominantly in Eastern and Southern regions. To reliably apply a robust modelling approach to a Europe-wide context targeted data collection is required.

## Introduction

Multi-host wildlife diseases are associated with over 60% of human diseases and over 75% of livestock diseases (Cleaveland et al. 2001). Understanding the risks of such diseases and how to manage them, relies on an understanding of the more widespread and common wildlife species, since these will be the main drivers of disease maintenance. Where are these species, and how abundant are they? We might expect common species to be well surveyed, but the empirical evidence suggests otherwise, with common UK species such as the mole (*Talpa europaea*), field vole (*Microtus agrestis*), and rabbit (*Oryctolagus cuniculus*) having very large uncertainties in population size when a purely systematic approach to the data is used (Croft et al. 2017). Since the UK is relatively well surveyed for mammalian biodiversity, we would expect the European situation to be even more poorly defined. Recently, the European Food Safety Authority (EFSA) has funded extensive work of determining the distribution of wild boar (*Sus scrofa*) (ENETWILD-consortium et al. 2021; ENETWILD-consortium et al. 2019a) and other ungulates (ENETWILD-consortium et al. 2022b) and carnivores (ENETWILD-consortium et al. 2023b) through the ENETWILD project. Along with this, additional monitoring of mammals through *ad hoc* sightings and systematic density estimates have been produced for larger mammals (ENETWILD-consortium et al. 2023a; Smith et al. 2023) with the launch of a European Observatory of Wildlife (ENETWILD-consortium et al. 2022a).

Species distribution models (SDMs) based on sightings have become increasingly popular in ecology to support decision-making on issues of wildlife disease and conservation. Large data repositories such as the Global Biodiversity Information Facility (GBIF) and initiatives like European Observatory of Wildlife (EOW) together with accessible tools built in platforms such as R (R Core Team 2024) have driven this widespread adoption. Many different approaches have been proposed each with distinct advantages and disadvantages depending on the type of data that is available (Guillera-Arroita et al. 2015; Golding et al. 2018). Throughout the course of the ENETWILD project several approaches have been considered based on increasing degrees of assumption and complexity to account for biases such as uneven survey effort (e.g., ENETWILD-consortium et al. 2019a; ENETWILD-consortium et al. 2019b; ENETWILD-consortium et al. 2020; ENETWILD-consortium et al. 2021; ENETWILD-consortium et al. 2023b).

Initial models were based purely on presence data (ENETWILD-consortium et al. 2019a), often referred to as profile or presence-only models. Such models (e.g., BIOCLIM, DOMAIN, Mahalanobis distance, Isolation Forests) have been shown to perform reasonably when survey effort is unbiased, or when there no knowledge of survey effort or broad species range (i.e., when other models are required to use uninformed pseudo-data to enable fitting) (Song & Estes 2023). However, in practice this is rarely the case, and more complex models are generally favoured (Soley-Guardia et al. 2024). Like many, we progressed to consider a presence-background approach (MaxEnt) using presences of other “associated” species (target-group; Phillips et al. 2009) to define a background reflecting survey effort and delimiting this using available expert-drawn range maps to exclude areas where presence may be limited by dispersal (e.g., Hattab et al. 2017; Koch 2021). This produced more plausible maps but without information on true absences, and consequently prevalence, could only provide a measure of relative suitability (Golding et al. 2018) and were impossible to benchmark performance against the real-world in any meaningful way (Merow et al. 2013). To combat this issue the project introduced the possibility of quantifying survey effort (ENETWILD-consortium et al. 2019b; ENETWILD-consortium et al. 2020) and use this to explicitly model the observation and ecological processes as part of both a coupled Bayesian (ENETWILD-consortium et al. 2021) and sequential frequentist framework (ENETWILD-consortium et al. 2022b). While conceptually the former presented the most complete and elegant approach, it proved computationally too expensive to run at the geographic scales required and accounting for spatial dependencies, using the “off-the-shelf” tools available (Vieilledent et al. 2014). The project instead pursued the latter approach, first estimating observability to identify “well-surveyed” locations (e.g., Lobo et al. 2018; Hill 2012) and then using the resulting presence-absence data to model the ecological process. This approach opens up a wider range of available modelling approaches including powerful machine-learning algorithm such as decision trees (Elith et al. 2008) and random forests (Valavi et al. 2021) which have been shown to consistently perform well against other models (e.g., Croft et al. 2017).

Despite every effort, correlative SDMs are never perfect, and performance is known to become unreliable when extrapolating beyond the limits of training data (Briscoe et al. 2019). As such it is always recommended to accompany predictions with an assessment of transferability such as a Multivariate Environmental Similarity Surface (MESS) (Elith et al.2010). This defines areas where the model projection is very poor due to, for example, local environmental or climatic factors that are not well represented in the training data. This could suggest areas where further data collection would be most informative to the collection of SDMs. Indeed, only a small degree of targeted sampling seems to be sufficient to substantially improve SDMs (Mondain-Monval et al. 2024).

Biodiversity data in Europe is very biased toward northern and western countries (García-Roselló et al. 2023) and such spatial bias can substantially impact model output (Soley-Guardia et al. 2024). The aim of this work is to use a generic species distribution modelling approach accounting for this bias to generate predicted distributions for common and widespread European mammals that are potentially involved in diseases of human or livestock concern. Areas where the models perform poorly can then be used to define geographic areas (countries) where future data collection would be most informative for improving such models, essential for improving future schemes of biodiversity monitoring in Europe (Liquete et al. 2024).

## Methods

### Overview

Our model framework comprised of three distinct steps (Fig. 1): (i) determine the long-term “stable” species range within which occurrence (presence and absence) could be assumed a consequence of environment rather than other factors such as dispersal limitation (delimitation) or recent incursion/retreat (non-stationarity); (ii) estimate the relative detectability/reportability of the species and by extension identify “well-surveyed” sites where frequency of observation may begin to reflect density (sufficient repeat visits to infer absence, or zero-density); (iii) assess habitat preferences and predict suitability across Europe. We applied this framework for twelve common mammal species (eight medium/large mammals: four ungulates, two lagomorphs, a medium-size rodent and a carnivore; two small rodents; and two bat species) that have a key involvement in various diseases of concern within Europe, posing a threat to the economy, livestock, or human health (e.g., African swine fever, Lyssaviruses including classical rabies, vector-borne diseases, and parasitic infections such as *Echinococcus multilocularis*) with the view that the resulting suitability maps may be used to inform future risk assessment and contingency planning.

**Figure 1:**
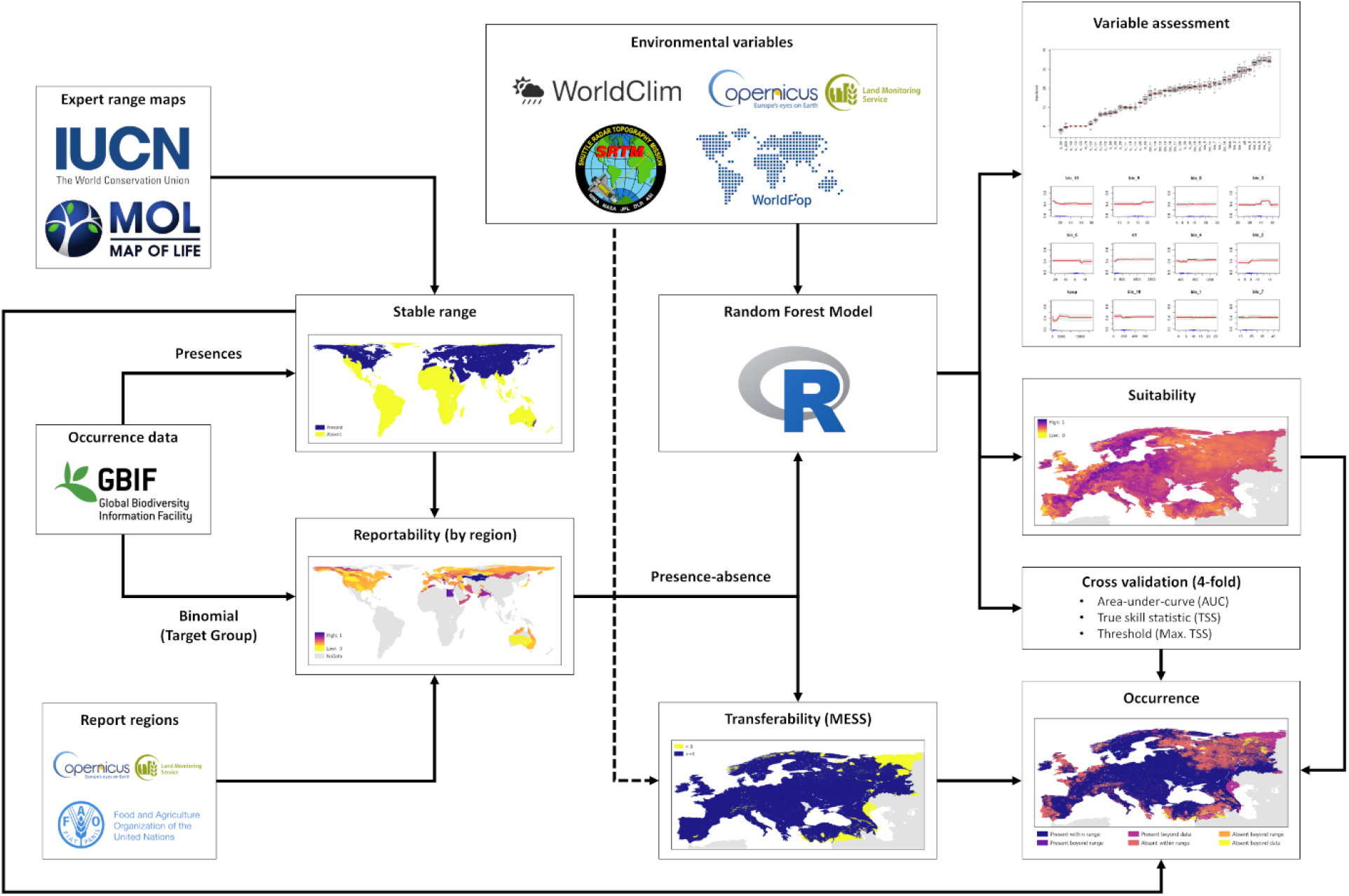
Schematic of model framework. Opportunistic occurrence data is first processed to generate a strictly presence-absence dataset where populations are considered stable before being applied in a random forest model to predict habitat suitability across Europe. In each element of the framework logos represent the sources of resources used. Occurrence data was obtained from GBIF (https://www.gbif.org); expert-drawn range maps were combined from IUCN (https://www.iucnredlist.org) and Map of Life (https://mol.org); reporting regions were defined based on biome derived from Copernicus GLS (https://zenodo.org) and administrative boundaries from the FAO GAUL dataset (https://data.apps.fao.org); environmental variables representing climate, land cover, topography, and human population were taken from Worldclim (https://www.worldclim.org), Copernicus GLS (https://land.copernicus.eu/global/products/lc), NASA/USGS SRTM (https://www.earthdata.nasa.gov), and Worldpop (https://www.worldpop.org), respectively. Random forest modelling was carried out using the “extendedForest” package in R (https://rdrr.io/rforge/extendedForest) with support to assess model performance from the “dismo”, “PresenceAbsence”, and “pdp” packages (https://cran.r-project.org).

### Stability

To estimate a “stable” range we supplemented information from expert drawn maps, typically representing long-term (30-year) broad-scale presence (within 50-100 km), with available occurrence data. Following the approach used by Map of Life (https://mol.org), we collated expert maps from multiple sources (Burgin et al. 2020, IUCN 2021, MDD 2020, Wilson et al. 2009-2019) and aggregated them to produce a single consensus (union) which we assumed to reflect the 30-years from 01/01/1991 to 31/12/2020. Sightings records (globally) for this period were obtained from GBIF (https://doi.org/10.15468/dl.rpr34v) and filtered to exclude any without an exact taxonomic description to species level or a coordinate uncertainty greater than 10 km. The resulting dataset was processed to describe presence for the species of interest in terms of location, centroid of a 10×10 km grid cell (EASE 2.0 spatial projection), and year of observation.

To establish evidence of consistent occurrence we partitioned presence data into a series of rolling 10-year windows (21 in total across the 30-year time-period). For each window, we used an approach combining local convex hulls (LoCoH; Maes et al. 2015) to enclose presences (including vertices representing the expert drawn boundary) assuming a maximum neighbour distance (defining relative locality) of 50 km. The stable range (which inherently included the complete expert derived map) was the intersection of ranges across all time windows, i.e., regions where there was consistent evidence of species presence (average value/probability of presence across windows equal to 1). To account for the uncertainty around presences (limited to 10 km accuracy) the final range was rasterized to a 10 km resolution grid before being disaggregated to a 1 km resolution and masked to only include land surface (see Appendices S1-S12).

### Reportability

To assess the reportability of a species, and by extension derive a strictly presence-absence dataset (i.e., one where there has been sufficient effort to declare a species present or absent), we considered sightings data (obtained from GBIF; https://doi.org/10.15468/dl.rpr34v) collected globally over the most recent 10-year period (01/01/2011 to 31/12/2020) using a mixture of different studies and protocols including systematic scientific studies but most commonly from citizen science driven recording apps. Using these records, we derived a binomial dataset describing distinct visits (dates) to a site (1×1 km grid cell) where either our target species had been reported (detections) or where it had not but an “associated” species (based on survey method: mid/large - visual, small - trap and bat - audio; Croft & Smith 2019, Coomber et al. 2021) had been (non-detections); the combination of which provided an estimate of total survey effort (Phillips & Dudík 2008). A list of species associations/groupings is provided in Appendices (S0). Sites where only the target species had been reported (presence-only) were excluded. Before continuing, we further filtered this dataset to remove sites where populations were thought to be unstable (beyond the estimated “stable” range).

To account for potential local differences which may influence the relative reporting rate of the target species compared with other associated species (e.g., culture/society/legislation and ecology/abundance/richness) we performed assessment on a regional basis delineated by distinct combinations (intersection) of country (administrative - reporting behaviour/importance; FAO 2015) and biome (community – species richness/abundance; Buchhorn 2022a). For each region, we computed the average reporting probability (total detections divided by total visits) across known presence locations (at least one detection) iteratively increasing the minimum levels of survey effort (visits) required by sample sites (presences) until the type II error reached zero (i.e., no possibility of exclusion of any locations where the species had zero detections but was in fact present). The reciprocal of this reporting probability, representing the expected number of visits for a species to be reported if present, was taken as a minimum threshold to declare absence and by extension consider a site “well-surveyed”. To maintain statistical rigor, we required effort thresholds to be estimated based on a sample size of at least 30 locations. Finally, applying thresholds to all locations (including those with presence data) on a region-by-region basis we generated a strictly presence-absence dataset.

### Suitability

Using experience from previous work comparing different models (Croft et al. 2017, Valavi et al. 2021), we opted to fit a random forest regression model (at global extent) considering variables describing climate, land use, topography, and human disturbance (Table 1) to explain our derived presence-absence data. While predictions from tree-based approaches such as random forest models are not overly sensitive to correlation amongst variables (Elith et al. 2008), assessment of variable importance and partial dependence which are based on permutation can be confounded. To overcome this issue without pre-selecting variables to retain we applied the approach outlined by Smith et al. (2011) using the “extendedForest” package in R, conditioning branching in a random forest model to only retain one variable per cluster based on pairwise correlations (“corr.threshold”) of 0.5 or greater. Following recommendation, we set variables “maxLevel” based on our sample size (to ensure a minimum of two per partition), “mtry” equal to the number of variables (Strobl et al. 2008), and “ntree” to 100. For each fit, we performed ten repetitions (1000 trees in total) and aggregated the results to produce plots of variable importance and partial dependence (figures 7 and 8 in Appendices S1-S12) and predict likelihood of occurrence (habitat suitability) across a European extent (Fig. 4 in S1-S12). At this stage spatial autocorrelation/dependence within model residuals was assessed by partitioning neighbour interactions into 5km wide bins and computing the permutation-based Pearson’s correlation coefficient (using the “permcor” from the R package “Rfast”) and corresponding p-value to test for significance, to determine if further action was required. Significant spatial dependence can be mitigated by spatially thinning training data (see spThin; Aiello-Lammens et al. 2015) before repeating the fitting process. This process is stochastic in nature and so if applied requires multiple repetitions. Here, we would suggest ten repetitions to be reasonable to represent sufficient variation without exceeding computational limits.

**Table 1:** List of model variables describing climate, land cover, topography, and anthropogenic influence used to assess habitat suitability.

| Code | Variable description | Code | Variable description |
| --- | --- | --- | --- |
| Climate<sup>1</sup> |  | Landcover<sup>2</sup> |  |
| bio1 | Annual mean temperature | lc111 | Closed forest (>70%), evergreen needle leaf |
| bio2 | Mean diurnal range | lc112 | Closed forest, evergreen broad leaf |
| bio3 | Isothermality | lc113 | Closed forest, deciduous needle leaf |
| bio4 | Temperature seasonality | lc114 | Closed forest, deciduous broad leaf |
| bio5 | Max temperature of warmest month | lc115 | Closed forest, mixed |
| bio6 | Min temperature of coldest month | lc116 | Closed forest, unknown |
| bio7 | Temperature annual range | lc121 | Open forest (15-70%), evergreen needle leaf |
| bio8 | Mean temperature of the wettest quarter | lc122 | Open forest, evergreen broad leaf |
| bio9 | Mean temperature of the driest quarter | lc123 | Open forest, deciduous needle leaf |
| bio10 | Mean temperature of warmest quarter | lc124 | Open forest, deciduous broad leaf |
| bio11 | Mean temperature of coldest quarter | lc125 | Open forest, mixed |
| bio12 | Annual precipitation | lc126 | Open forest, unknown |
| bio13 | Precipitation of wettest month | lc20 | Shrubs |
| bio14 | Precipitation of driest month | lc30 | Herbaceous vegetation |
| bio15 | Precipitation seasonality | lc40 | Cropland |
| bio16 | Precipitation of wettest quarter | lc50 | Urban/built-up |
| bio17 | Precipitation of driest quarter | lc60 | Bare / sparse vegetation |
| bio18 | Precipitation of warmest quarter | lc70 | Snow and Ice |
| bio19 | Precipitation of coldest quarter | lc80 | Permanent water bodies |
| Anthropogenic/Topographic |  | lc90 | Herbaceous wetland |
| hpop | Human population density<sup>3</sup> | lc100 | Moss and lichen |
| alt | Mean altitude<sup>4</sup> | lc200 | Open seas |

<sup>1</sup> Derived from Worldclim historic monthly weather data 2010-2019, CRU-TS 4.06 (Harris et al. 2020) downscaled with WorldClim 2.1 (Fick & Hijmans 2017), using the “biovars” function from the R “dismo” package (Hijmans et al. 2023).
<sup>2</sup> Derived from Copernicus Global Land Service: land cover 100m collection 3 2019 (Buchhorn et al. 2020b).
<sup>3</sup> Derived from Worldpop global population density 2015 (https://www.worldpop.org).
<sup>4</sup> Derived from USGS Space Shuttle Radar Topography Mission (SRTM) GL30 (https://lta.cr.usgs.gov/SRTM1Arc).

To evaluate the transferability of our model predictions, we performed a MESS analysis (Elith et al., 2010) which identifies regions whose environmental conditions are deemed insufficiently represented by the training dataset and so may produce unreliable predictions. Finally, model performance was assessed using 4-fold cross validation with filtering to remove spatial sorting bias (Hijmans 2012) to compute several common metrics for predictive accuracy: AUC (area under curve statistic, calibrated against a null model; Hijmans 2012); TPR (True positive rate - Sensitivity); TNR (True negative rate - Specificity); and TSS (True skill statistic) based on an optimal threshold value (THD) to maximise TSS (Liu et al. 2013). Partitioning for cross validation was done in two different ways: (i) unbiased random selection producing environmentally similar training and testing datasets; (ii) selection according to environmentally clustering producing distinct training and testing datasets (assessed using a MESS analysis). Evaluation under both routines is important to provide insights into the predictive performance of models within and beyond the limits of available training data respectively (Roberts et al. 2017).

## Results

While for some species there was evidence of significant spatial autocorrelation in residuals compared with a random sample, correlation coefficients for close distances (immediately adjacent; 1 km) were low (<0.05) and as such removal using available methods (thinning) was considered to provide limited benefit for the additional computational cost (balance of presence-absence data shown in Table 2).

**Table 2:** Summary of available presences and assumed background/absences (sampled 1 km raster cells) for each species across a global extent. Background refers to cells with at least one inferred visit where the target species could have been observed. Counts are for cells within the assumed “stable” species range only. Values in brackets denote proportion of cells where species presence was reported.

| Specie | Range | Presence-Background | Presence-Absence |
| --- | --- | --- | --- |
| *Alces alces* | 28,729,250 | 77,947 (0.075) | 4,814 (0.3) |
| *Capreolus capreolus* | 7,361,908 | 195,813 (0.144) | 37,087 (0.419) |
| *Cervus elaphus* | 6,369,489 | 197,837 (0.024) | 24,615 (0.069) |
| *Eptesicus serotinus* | 13,199,216 | 20,019 (0.159) | 3,069 (0.386) |
| *Lepus europaeus* | 11,778,450 | 163,316 (0.092) | 22,954 (0.303) |
| *Microtus agrestis* | 12,219,968 | 16,528 (0.066) | 1,597 (0.221) |
| *Myodes glareolus* | 9,121,415 | 16,324 (0.11) | 1,609 (0.387) |
| *Myotis daubentonii* | 8,200,392 | 26,664 (0.132) | 4,509 (0.329) |
| *Oryctolagus cuniculus* | 3,150,771 | 189,374 (0.106) | 32,635 (0.31) |
| *Sciurus vulgaris* | 24,486,166 | 193,391 (0.104) | 52,190 (0.291) |
| *Sus scrofa* | 30,182,163 | 115,077 (0.065) | 6,643 (0.345) |
| *Vulpes vulpes* | 64,653,552 | 317,524 (0.094) | 25,556 (0.386) |

In general, the predictive performance of models within the environmental limits of the training data (random fold selection for cross-validation - interpolation) across all species was good/acceptable based on AUC showing a score of 0.7 or greater except for *Myodes glareolus* which only achieved an AUC of 0.66 (Table 3). However, when cross-validation was performed deliberately partitioning folds based on environmental similarity (thereby evaluating the ability of models to predict beyond the available training data - extrapolation) evaluation metric decreased substantially with none remaining above the 0.7 threshold generally accepted to indicate moderate to good performance (Table 3). This is expected for non-parametric machine learning approaches like random forest where there is no information to underpin response beyond given variable limits. Evaluation based on TSS potentially indicated a lower degree of predictive performance, although expectations of what constitutes acceptable performance is less well defined. A score of 0.4, like that applied to the kappa statistic (Landis & Koch 1977), has previously been proposed. By this benchmark only predictions for elk (*Alces alces*), roe deer (*Capreolus capreolus*) and brown hare (*Lepus europaeus*) would be considered reasonable. Like AUC, performance based on TSS was notably reduced when extrapolating to new environmental conditions beyond the scope of the model training data.

**Table 3:** Summary of model predictive performance for each species based on a 4-fold cross validation with unbiased partitioning (within the same environmental envelope as the training dataset) adjusted for spatial sorting bias (Hijmans 2012). Values in brackets denote results of a 4-fold cross validation with environmentally clustered partitioning to illustrate performance beyond the environmental envelope of the training dataset. For each species, scores provide the mean (across folds) area under curve statistic (AUC), the true-skill statistic (TSS) with corresponding threshold (THD), sensitivity (SE) and specificity (SP), and the proportion of the testing data that is beyond the environmental envelope bounding the training data (MESS).

| Specie | AUC | TSS | THD | SE | SP | MESS |
| --- | --- | --- | --- | --- | --- | --- |
| *Alces alces* | 0.78 (0.62) | 0.43 (0.22) | 0.36 (0.42) | 0.77 (0.73) | 0.66 (0.49) | 0.01 (0.57) |
| *Capreolus capreolus* | 0.80 (0.62) | 0.47 (0.18) | 0.45 (0.43) | 0.79 (0.74) | 0.68 (0.45) | 0.00 (0.08) |
| *Cervus elaphus* | 0.74 (0.66) | 0.35 (0.24) | 0.22 (0.17) | 0.66 (0.82) | 0.69 (0.42) | 0.00 (0.31) |
| *Eptesicus serotinus* | 0.72 (0.53) | 0.36 (0.06) | 0.42 (0.43) | 0.69 (0.66) | 0.67 (0.41) | 0.02 (0.54) |
| *Lepus europaeus* | 0.80 (0.61) | 0.50 (0.16) | 0.33 (0.42) | 0.81 (0.59) | 0.69 (0.58) | 0.00 (0.20) |
| *Microtus agrestis* | 0.70 (0.65) | 0.33 (0.27) | 0.33 (0.38) | 0.64 (0.80) | 0.70 (0.47) | 0.02 (0.58) |
| *Myodes glareolus* | 0.66 (0.55) | 0.26 (0.12) | 0.42 (0.54) | 0.59 (0.46) | 0.66 (0.67) | 0.03 (0.43) |
| *Myotis daubentonii* | 0.70 (0.58) | 0.32 (0.11) | 0.38 (0.28) | 0.64 (0.90) | 0.68 (0.20) | 0.01 (0.55) |
| *Oryctolagus cuniculus* | 0.75 (0.59) | 0.38 (0.15) | 0.37 (0.43) | 0.68 (0.70) | 0.70 (0.45) | 0.00 (0.30) |
| *Sciurus vulgaris* | 0.73 (0.65) | 0.33 (0.22) | 0.43 (0.48) | 0.69 (0.57) | 0.64 (0.65) | 0.00 (0.37) |
| *Sus scrofa* | 0.74 (0.54) | 0.37 (0.08) | 0.33 (0.42) | 0.78 (0.76) | 0.59 (0.32) | 0.00 (0.59) |
| *Vulpes vulpes* | 0.74 (0.60) | 0.36 (0.17) | 0.39 (0.46) | 0.73 (0.55) | 0.64 (0.62) | 0.00 (0.22) |

Evaluation of variable importance suggested land cover was broadly less important than other factors such as climate (e.g. bio15), altitude (alt), and human population (hpop) (Fig. 2). However, this is likely a reflection of high levels of co-correlation between these variables and therefore does not necessarily reflect their individual importance (that all are equally more important than land cover) but that of the collective. Land cover variables are not highly correlated and highlight various species preferences matching expectation. For example, elk, red squirrel (*Sciurus vulgaris*), and red deer (*Cervus elaphus*) all show a high variable importance for forest cover (lc_1XX codes), brown hare (*Lepus europaeus*) shows high importance for cropland cover (lc40), roe deer and European rabbit (*Oryctolagus cuniculus*) shows high importance for human population (hpop), and bank vole (*Myodes glareolus*) and Daubenton’s bat (*Myotis daubentonii*) show high importance for water bodies (lakes and rivers; lc_80). Other species, in particular, red fox (*Vulpes vulpes*) or serotine bat (*Eptesicus serotinous*) show less pronounced association reflective of a more generalist species. Examining variable importance for wild boar (*Sus scrofa*) in more detail, our results suggest altitude, forest, and urban land cover as being most influential followed by climatic variables relating to precipitation in the warmest quarter (bio18) and temperature in the wettest quarter (bio8), which for the UK at least typically corresponds to winter, and the precipitation in the warmest quarter (bio18) which reflects the likelihood of drought conditions. This shows good agreement with previous model studies (ENETWILD-consortium et al. 2018).

**Figure 2:**
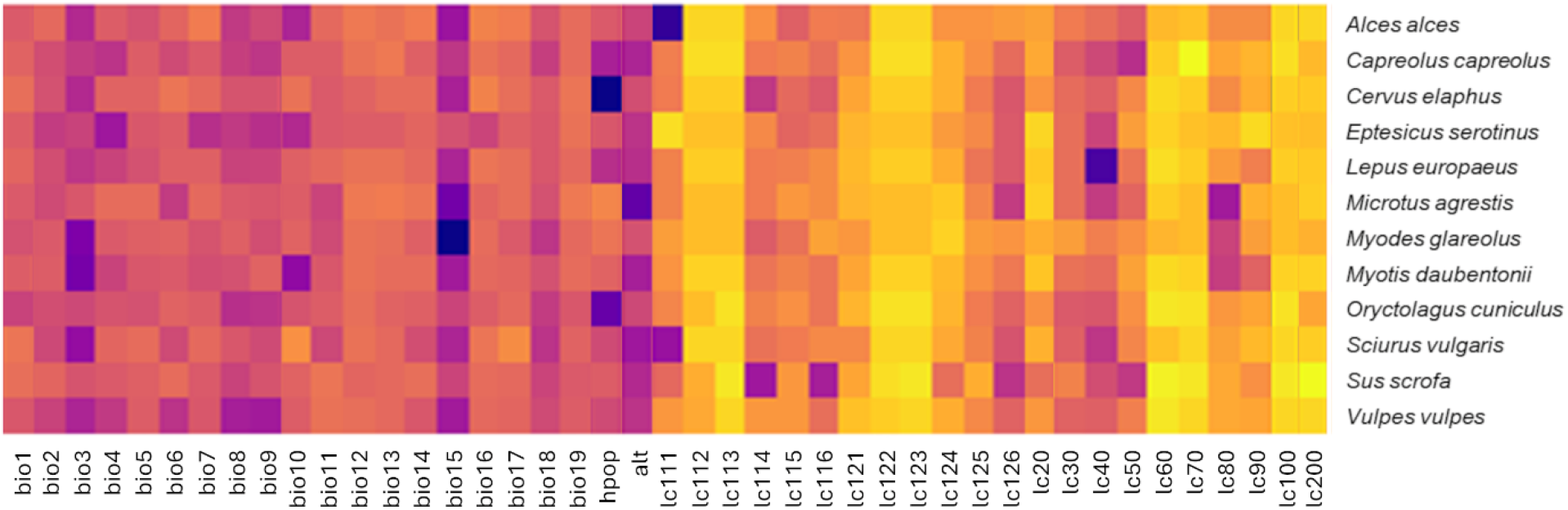
Summary of variable importance across species. Darker (more intense) colours reflect greater normalised (0-1) permutation importance. Overall, as a group land cover variables are notably less important. However, this is most likely a reflection of the correlation between other variables, particularly topography and climate variables, which can make the precise contribution of each difficult to distinguish (i.e., if one is considered important, based on the data available, other highly correlated variables will appear similarly important). Specific land cover variables feature prominently for species with known specialism/dependence (e.g., red squirrels, boar, and elk are associated with forest - lc1XX, brown hare with cropland - lc40, bank voles and Daubenton’s bats with permanent water bodies - lc80).

For each species, the result of multi-variate environmental similarity surface (MESS) analyses was clipped to exclude cells without evidence of species presence (i.e., strong evidence of absence due to ecological conditions well beyond species range, rather than a lack of survey effort) using derived range maps (union across rolling windows i.e., a value/probability of presence >0). The resulting maps were combined across all species (sum per cell of values less than 0; Fig. 3) and aggregating by country (mean of sum across cells; Table 4) highlighted countries within Europe where targeted data collection may improve the transferability of model predictions. In general, data gaps in Eastern and Southern European countries (Russia, Ukraine, Turkey, Romania etc) are widespread and would benefit from a country-wide effort whereas data gaps in Northern and Western countries (Switzerland, Austria, Andorra, Norway) are more localised (e.g., areas of the Alpes and Pyrenees).

**Table 4:**
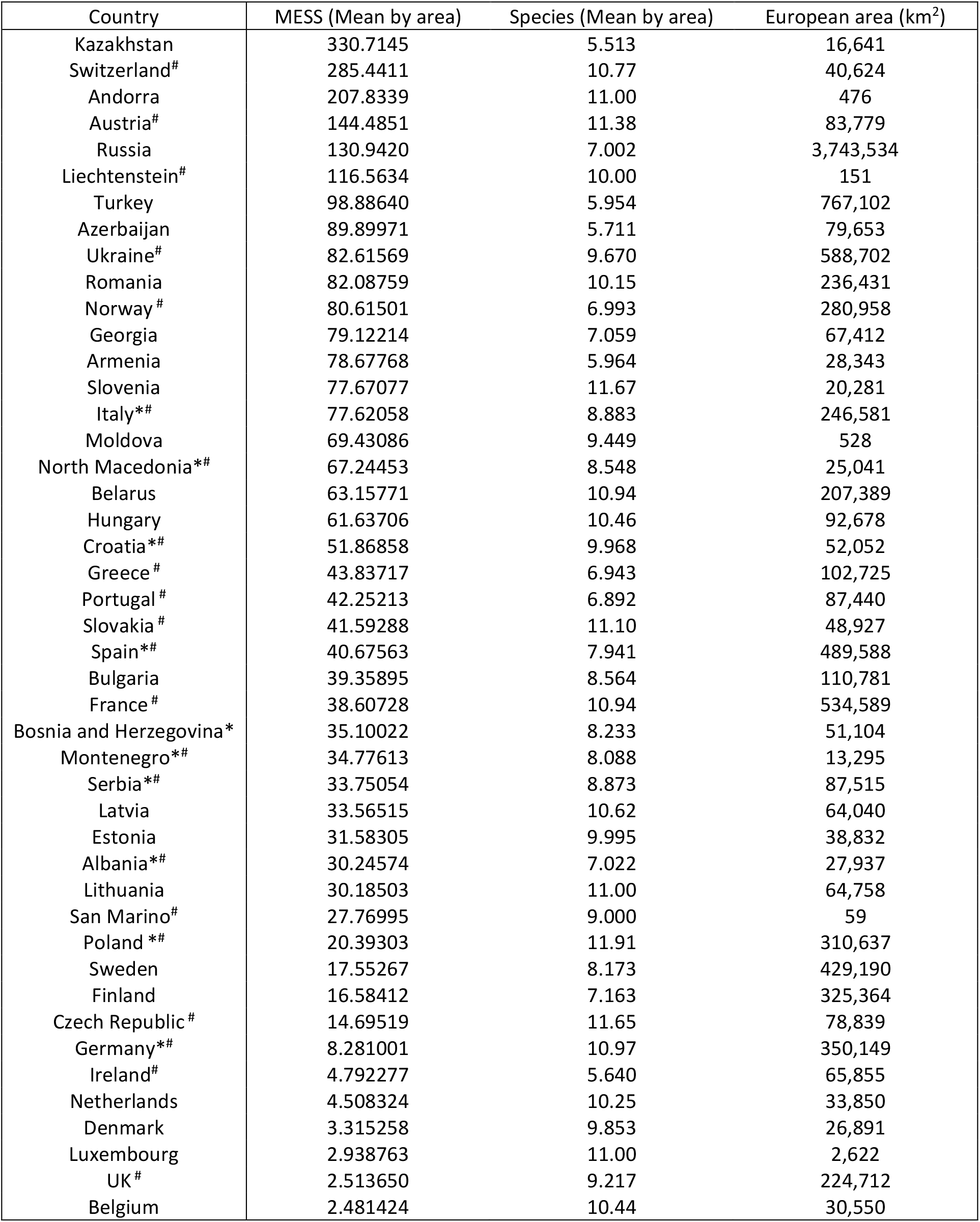
Summary of environmental similarity to available data aggregated across species by country. Order indicates possible priority for targeted data collection to improve model transferability (highest first). Countries marked with an asterisk (*) denote those where the ENETWILD-MammalNet tools (https://mammalnet.com) has been launched. Those marked with a hash (^#^) denote countries whose predominant language is available in the iMammalia app.

**Figure 3:**
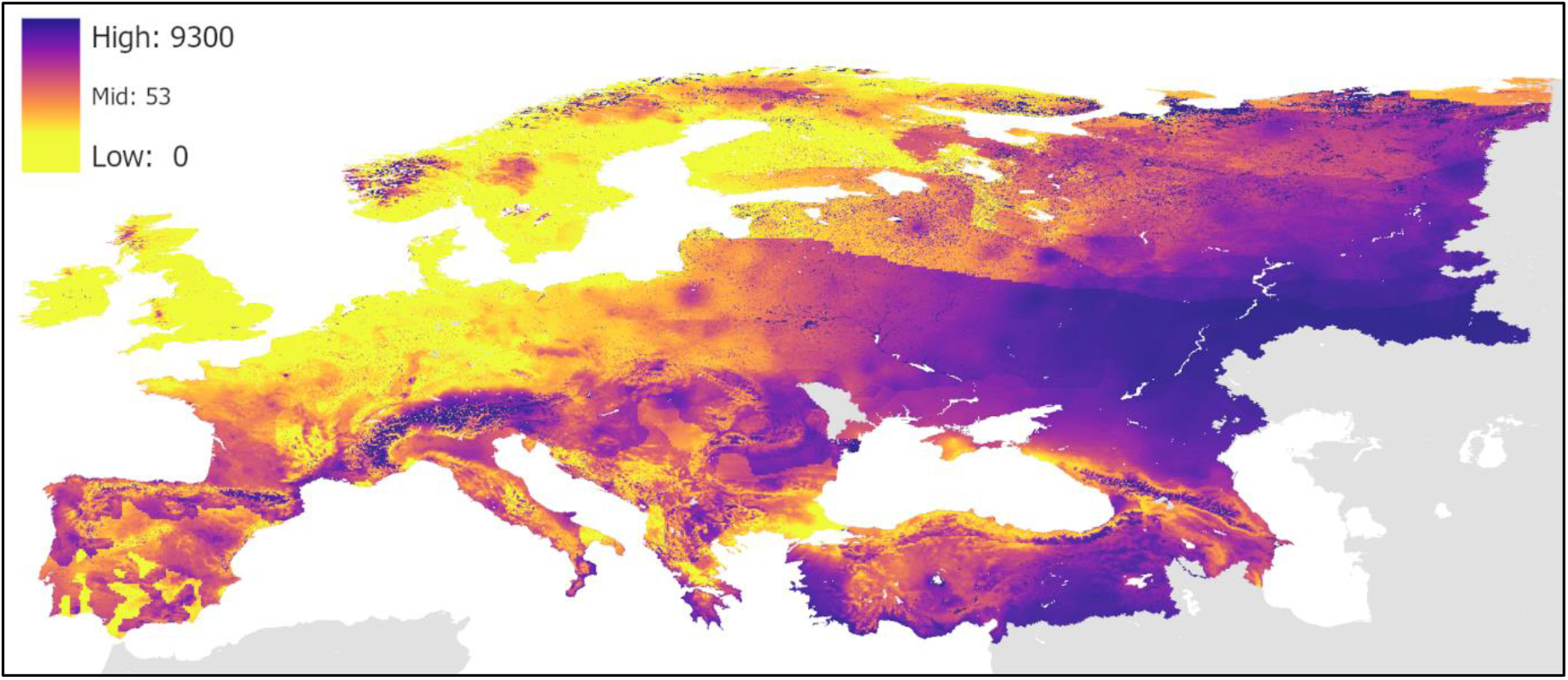
Map showing the novelty of environmental conditions compared to current available training data combined across all twelve species considered in this study (aggregated MESS analysis outputs delimited to exclude regions beyond evidence of species occurrence). Darker (more intense) colours represent greater novelty and by extension we suggest a higher priority for targeted data collection. Northern and Western Europe is reasonably well represented by current data with only some localised patches that would benefit from data collection. Eastern and Southern Europe would benefit more widely from targeted data collection from programs such as EOW.

## Discussion

Mammalian biodiversity data recording is now very widespread with a diverse range of actors involved. In 2023 alone GBIF reported over 600,000 mammal records in Europe. From a European perspective we could then ask where would additional recording effort be best placed? Here we examined this question for common species most likely to be involved in multi-host diseases of concern to humans and livestock. There may be a different output metric if we were concerned about species conservation, or invasive species, but, if we cannot even describe the common species distribution and abundance well, then the bias in sampling effort is highly likely to lead to false conclusions.

We report here on the long-term development of a generic framework to predict habitat suitability based on opportunistic occurrence data for common widespread European mammals, since of all species these should be the most reliable. We use occurrence data as these are available for all mammal species, thus ensuring a consistent approach, whereas the use of, say, hunting data would limit this approach to species where hunting records are reported in high detail. The modelling approach performs well within the environmental limits of the available training data but as expected performance, and by extension predictions, become more unreliable beyond these limits. Given this limitation of modelling, it is perhaps surprising, and concerning, that such large geographical areas are identified by the MESS analysis across these twelve species (Fig. 3).

The models seem to perform worst, lower AUC and TSS, for the two bat and two vole species (Table 3). These species are mostly recorded through specific methods (audio or small mammal traps), with the small rodents among the least recorded mammals for their population size (Croft et al. 2017). The potential for acoustic monitoring of small terrestrial mammals and bats may lead to an increase in these records over time which may improve model performance (Newson & Pearce 2022). However, some refinement of the general modelling approach may also be beneficial such as considering a finer spatial scale more representative of the species ecology and consideration for species assemblages at a local scale, for instance to remove records of locally sensitive or rare species which may not fully represent survey effort for common species (Coomber et al. 2021).

Despite the data limitations, the proposed modelling method performs well across most of the species considered (mid-large mammals). Nevertheless, there are modifications to the framework, beyond those already mentioned specifically for small mammals and bats, which have potential to yield some improvement in performance and merit some further investigation. We make several necessary assumptions about the scale of variation in detectability. Where data allows, we could consider finer-scale delimitation in reporting practices to explore within-country variation such as county-level in the UK. Data from south Europe (in particular for Spain) shows large variation in recording intensity which may be indicative of different cultures/attitudes/policies across autonomous regions but also can be associate to bioregion effect (ENETWILD-consortium et al. 2020). Introduction of a calibration step could be considered to simplify models. Random forest models are not known to be especially vulnerable to overfitting however applying some degree of variable selection (e.g., VSURF; Genuer et al. 2015) may help to discount redundancy and by extension provide a clearer picture of model transferability (assuming redundant variables can be assumed to remain so when projecting into new environmental space). Applying downscaling random forests to balance classes (presence and absences) instead of regression can also be used to avoid overfitting (Valavi et al. 2021), reducing the impact of any overlap between presences and absences to produce shallower trees.

In some cases, we have noted the influence of spatial dependence (autocorrelation) in model residuals. While spatial thinning can be used to address this issue (e.g., Aiello-Lammens et al. 2015) for binary data this can be complicated (Steen et al. 2021), and care must be taken to avoid introducing error (Ten Caten & Dallas 2023, Lamboley & Fourcade 2024). Alternative methods which seek to represent spatial dependence directly within models such as SpatialRF (Hengl et al. 2018, Benito 2021) may provide a more promising solution. Our analysis here is based only on the highest quality data we can identify. We ignore any data that falls below this standard. However, this data is derived from the same underlying ecological system and may therefore be helpful to benchmark predictions. Integration of different methods based on lower standards of data (e.g., presence-background or even presence-only) may improve performance and potentially help to extend transferability. Similarly, consideration for existing ecological knowledge, perhaps to identify regions of unsuitability where species are expected to be absent as a counterpoint to stable presence, may also improve predictive performance.

These developments notwithstanding (based on available presence-absence data only), the MESS analysis identified areas in Europe where further mammal data collection would be most useful. This did not identify specific areas for specific species (see supplementary files), but rather areas where generic methods would be most beneficial. This includes ad hoc reporting (e.g., iMammalia: https://european-mammals.brc.ac.uk or iNaturalist: https://www.inaturalist.org/) or ad hoc camera trapping (e.g., MammalWeb: https://www.mammalweb.org/) as well as more systematic cameras trapping such as that being used in the European Observatory of Wildlife (EOW: https://wildlifeobservatory.org/). Additionally, it does not suggest that all areas identified in Figure 3 need to be surveyed, but that further records from each general habitat. Thus, surveys in Switzerland (the second highest priority country according to the MESS results) would reduce the importance of Andorra and Austria (listed third and fourth) as all these areas are mountainous. However, it is important to note that the MESS results need to be considered alongside how many species this relates to and the area. The MESS analysis may guide the identification of areas in Europe where further mammal data collection would be priority in the frame of future schemes of biodiversity at continental scale (Liquete et al. 2024).

This work indicates that despite the 600,000 mammal records reported in 2023 across Europe in GBIF, the geographical bias in data collection needs to be reduced to improve species distribution models for even the most widespread and common species. Evidence from the UK butterfly recording scheme indicates that even a 1% change in records toward areas identified by modelling would improve model output (Mondain-Monval et al. 2024). The next steps are therefore to promote data collection in specified areas and monitor improvement in the model output; combine model predictions with estimates of density (e.g., from EOW); and ensure the output is available to underpin risk assessment for relevant diseases.

## Supporting information

Recording assemblages

Output - Elk

Output - Roe deer

Output - Red deer

Output - Serotine bat

Output - European hare

Output - Field vole

Output - Bank vole

Output - Daubenton's bat

Output - European rabbit

Output - Red squirrel

Output - Wild boar

Output - Red fox

Occurrence data summary

## Supplementary material

Appendix S0 (.xlsx) list of assumed species associations / reporting assemblages.

Appendices S1-S12 (.pdf) contain separate model outputs for each individual species.

Appendix S13 (.pdf) table summarising data classifications (presence/absence, within/beyond range, within/beyond training data) for each species across Europe.

## Acknowledgements

This work was co-funded by the European Food Safety Authority (EFSA) as part of the ENETWILD project and the European Union’s Horizon Europe Project 101136346 EUPAHW. JA Blanco-Aguiar was funded by the European Commission - NextGenerationEU, through Momentum CSIC Programme (MNT24-IREC-01). We would like to acknowledge all of the organisations and individuals that have contributed to the datasets which make important work for public benefit such as this possible.

