## Supplementary material for "Predicting the distribution of common wild mammal species across Europe – are there sufficient occurrence data?": Output - Elk

### Appendix 1

Supplementary model results for ***Alces alces***

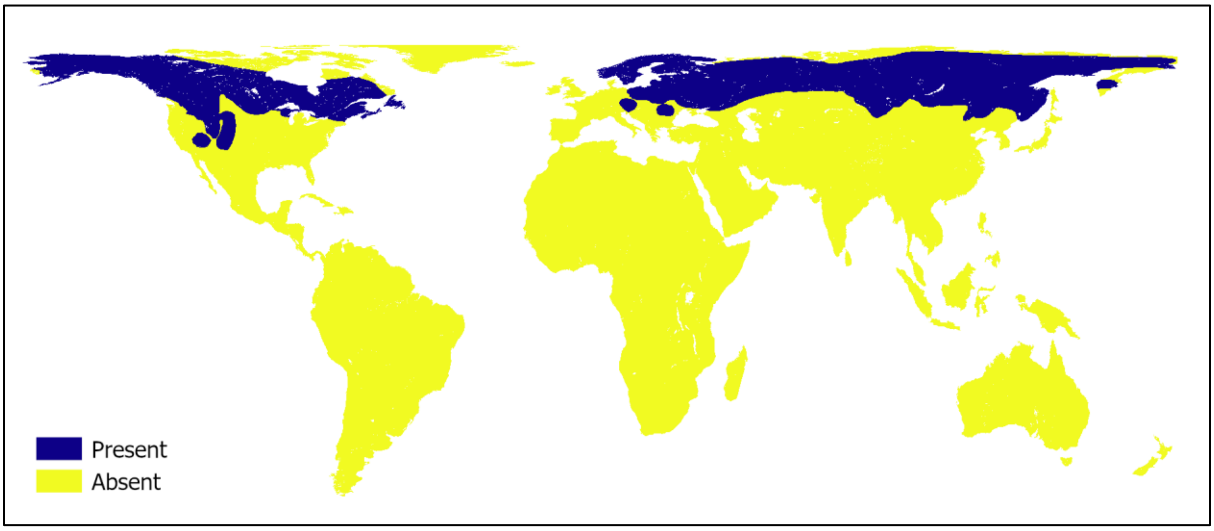


Figure 1: Map showing the assumed long-term "stable" range within which model training data is selected. Footprint combines occurrence data downloaded from GBIF (https://doi.org/10.15468/dl.rpr34v) with expert drawn maps from Map of Life (https://doi.org/10.48600/MOL-ZZRS-Q778; https://doi.org/10.48600/MOL-48VZ-P413; https://doi.org/10.48600/MOL-7R3J-8066) and IUCN (IUCN 2022. The IUCN Red List of Threatened Species. International Union for Conservation of Nature. Accessed 07/03/2023).

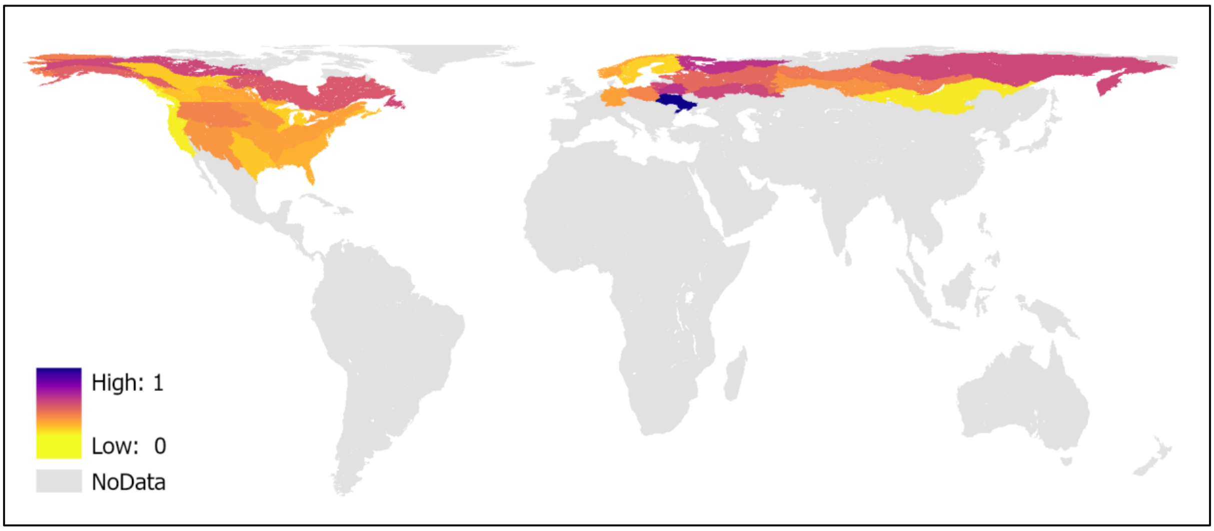


Figure 2: Map showing estimated detectability assuming presence calculated for regions reflecting a unique combination of landmass, bioregion (https://doi.org/10.5281/zenodo.5848610) and administrative (country level - GAUL0 https://data.apps.fao.org/map/gsrv/gsrv1/gaul/wms) boundaries.

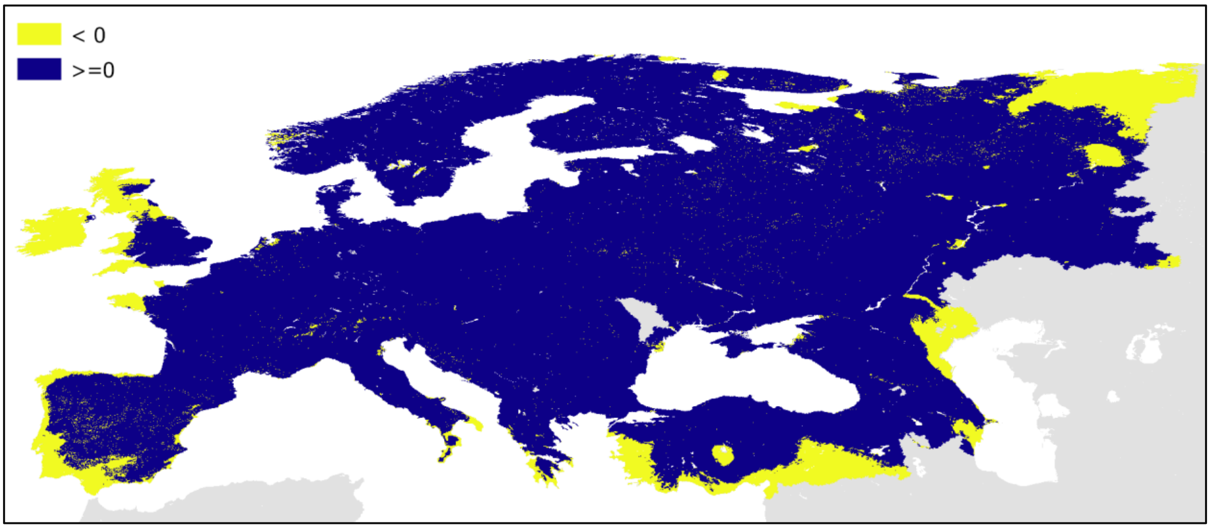


Figure 3: Map showing the transferability of model predictions (extent over which they can be considered reliable) based on a MESS analysis across a European extent. Positive values (>=0) denote environmental conditions represented by training data (inclusion). Negative values (<0) denote environmental conditions not represented by training data (omission) and therefore projections may be unreliable.

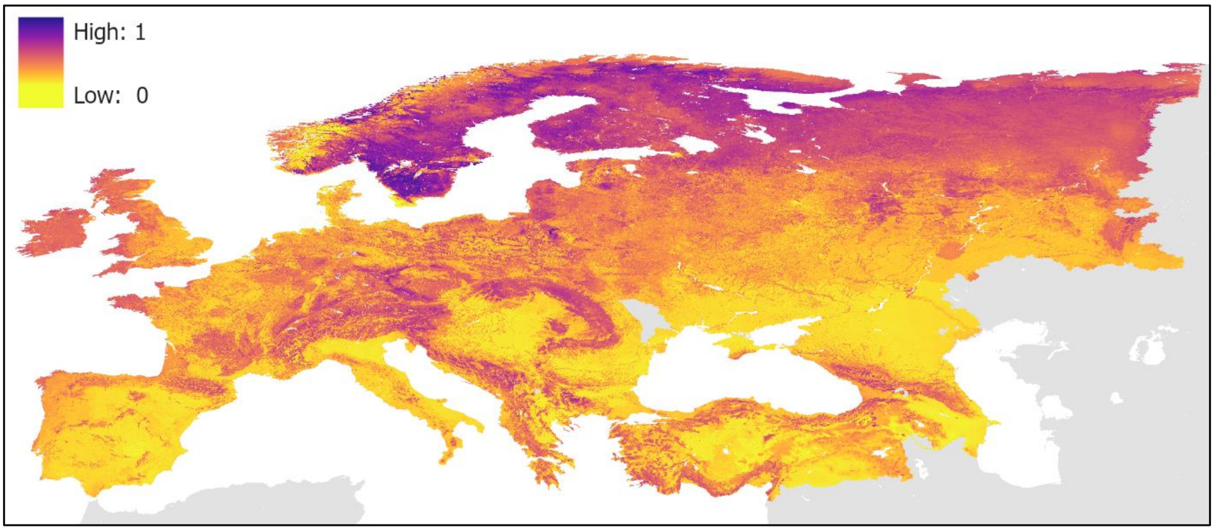


Figure 4: Map showing projected habitat suitability (probability of occurrence) across a European extent. Darker colours indicate greater suitability. Note map does not remove regions where prediction may be considered unreliable. This output should be used in conjunctions with the additional information presented in Figure 3.

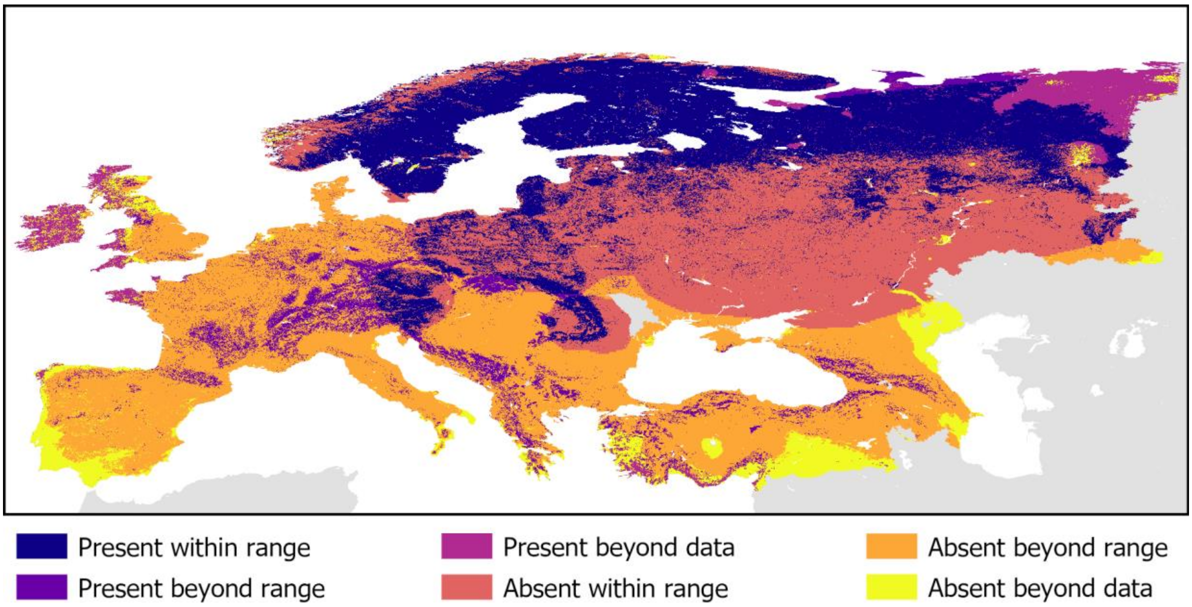


Figure 5: Map showing predicted occurrence across a European extent. Suitability threshold chosen to maximise the True Skill Statistic (TSS) measuring predictive performance averaged across estimates from a 4-fold cross validation. Colours denote combinations of presence/absence, within/beyond stable range and within/beyond the limitations of the training data (i.e., where predictions are reliable/unreliable).

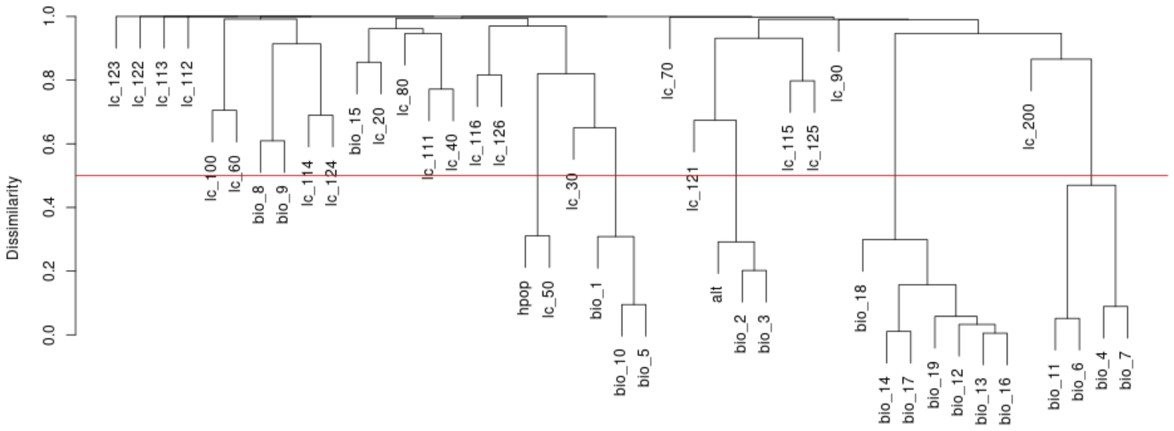


Figure 6: Hierarchical clustering (dendrogram) of variables based on distance using 1- Pearson's R (dissimilarity). Red line shows threshold of 0.3 below which variables are considered highly correlated.

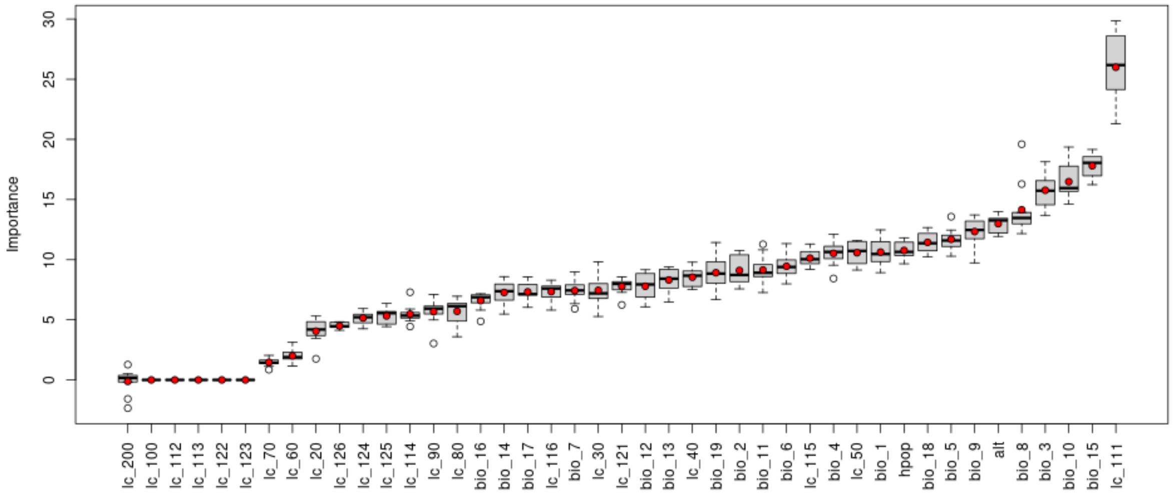


Figure 7: Permutation importance assessed across 10 stochastic replicates of model fit. Red dots denote mean importance across replicates.

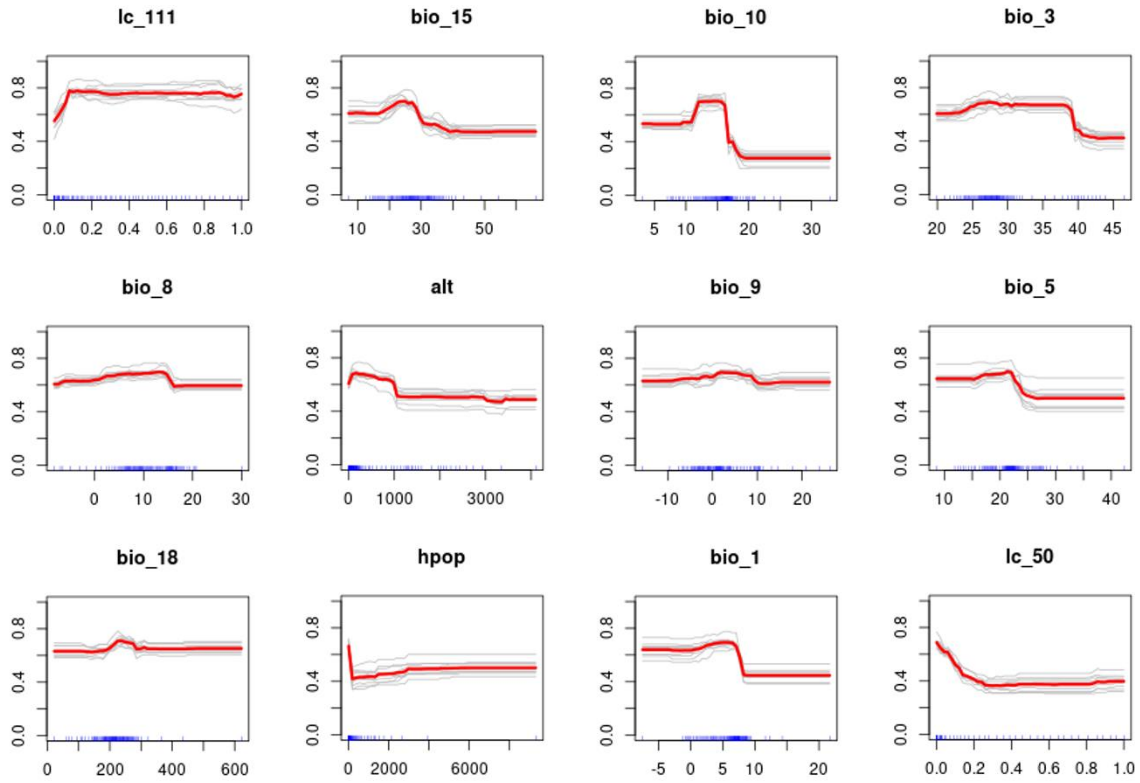


Figure 8: Partial dependence (response) plots for the twelve highest ranked (by importance) variables. Red lines denote mean response across 10 replicates. Blue ticks denote sample distribution (percentiles).
