## Supplementary material for "Predicting the distribution of common wild mammal species across Europe – are there sufficient occurrence data?": Occurrence data summary

**Table S1:** Summary of data classifications (present/absent, within/beyond range, within/beyond data) for each species across Europe. Numbers represent total area ($km^2$) for each classification (Appendices S1-S12 Figure 5).

| Specie | Present | | | Absent | | |
|---|---|---|---|---|---|---|
| | WR | BR | BD | WR | BR | BD |
| *Alces alces* | 2,694,891 | 554,004 | 484,230 | 2,756,971 | 3,112,437 | 589,601 |
| *Capreolus capreolus* | 4,451,221 | 241,018 | 1,175,035 | 2,236,597 | 873,270 | 1,214,993 |
| *Cervus elaphus* | 2,296,191 | 980,222 | 4,013,799 | 2,410,420 | 401,569 | 89,933 |
| *Eptesicus serotinus* | 559,977 | 9,167 | 6,972,967 | 690,368 | 95,508 | 1,842,250 |
| *Lepus europaeus* | 3,883,284 | 341,192 | 4,032,367 | 726,587 | 257,844 | 928,963 |
| *Microtus agrestis* | 1,505,999 | 44,630 | 8,240,940 | 253,317 | 32,819 | 92,532 |
| *Myodes glareolus* | 2,313,786 | 62,592 | 6,830,663 | 576,672 | 6,011 | 380,513 |
| *Myotis daubentonii* | 2,854,191 | 233,047 | 3,955,046 | 1,877,147 | 445,221 | 805,585 |
| *Oryctolagus cuniculus* | 532,434 | 256,580 | 1,485,294 | 1,868,447 | 882,958 | 5,144,525 |
| *Sciurus vulgaris* | 1,836,208 | 292,083 | 348,757 | 5,835,684 | 1,186,028 | 693,374 |
| *Sus scrofa* | 2,533,064 | 131,587 | 5,919,194 | 924,840 | 293,855 | 367,698 |
| *Vulpes vulpes* | 722,593 | 6,868,887 | 722,593 | 221,245 | 2,373,493 | 221,245 |

Within range (WR) | Beyond range (BR) | Beyond data (BD)
